# ICARUS v3, a massively scalable web server for single cell RNA-seq analysis of millions of cells

**DOI:** 10.1101/2023.11.20.567692

**Authors:** Andrew Jiang, Russell G Snell, Klaus Lehnert

## Abstract

In recent years, improvements in throughput of single cell RNA-seq have resulted in a significant increase in the number of cells profiled. The generation of single cell RNA-seq datasets comprising >1 million cells is becoming increasingly common, giving rise to demands for more efficient computational workflows. Here, we present an update to our single cell RNA-seq analysis web server application, ICARUS (available at https://launch.icarus-scrnaseq.cloud.edu.au/) that allows effective analysis of large-scale single cell RNA-seq datasets. ICARUS v3 utilises the geometric cell sketching method to subsample cells from the overall dataset for dimensionality reduction and clustering that can be then projected to the large dataset. We then extend this functionality to select a representative subset of cells for downstream data analysis applications including differential expression analysis, gene co-expression network construction, gene regulatory network construction, trajectory analysis, cell-cell communication inference and cell cluster associations to GWAS traits. We demonstrate analysis of single cell RNA-seq datasets using ICARUS v3 of 1.3 million cells completed within the hour.

## Introduction

With the increased throughput of single cell RNA-seq technologies in recent years, the necessity for large scale data analysis is becoming increasingly important. Single cell RNA-seq datasets and data from aggregated sources now include millions of cells (1,2) which has increased the need for efficient computational analysis. We have previously introduced ICARUS, an interactive web server application for single cell RNA-seq analysis (3,4). ICARUS utilises the Seurat R workflow to perform preprocessing, dimensionality reduction and clustering. Recently released Seurat v5 (https://cran.r-project.org/web/packages/Seurat/index.html) and the BPCells R package (https://github.com/bnprks/BPCells) introduces methods to store large datasets on-disk whilst utilising geometric sketch-based methods to identify a subpopulation of representative cells from the overall dataset to store in memory for rapid and iterative exploration. This drastically lowers computational processing time whilst retaining power to detect heterogeneity across the data. Our update to ICARUS v3 harnesses this methodology to perform dimensionality reduction and clustering as well as utilising this population of sketched cells to perform common downstream data analysis including co-expression network analysis (5), regulatory gene network construction (now updated to use the SCENIC+ regulatory motif database) (6), trajectory analysis (7), cell-cell signalling (8) and examination of cell cluster association with GWAS traits (3,4).

### Exceptional computational processing speed

ICARUS v3 implements the ‘geometric sketching’ method of sampling a subset of representative cells in the overall dataset. This method was first introduced by Berger and colleagues (9) and recently incorporated into the Seurat v5. Geometric sketching involves an approximation of the geometry of a single cell RNA-seq dataset by employing equal-volume boxes within multidimensional space that each cell occupies defined by its gene expression profile. These boxes are positioned to encompass all cells in the dataset, ensuring that each box contains at least one cell. Cells are then sampled at random from these boxes ensuring that both rare cell types and common cell types that occupy a similar volume of transcriptomic space are equally represented in the ‘sketched’ dataset (9). Once a subset of sketched cells is determined, this heavily reduced dataset is stored in memory while the larger overall dataset is stored on-disk using the BPCells R package (https://github.com/bnprks/BPCells). Dimensionality reduction and clustering are performed on the sketched dataset at efficient speed, and the resultant clusters from the sketched dataset are then projected back onto the overall dataset stored on disk (ProjectData functionality of Seurat v5). We demonstrate efficient clustering of a dataset comprised of 1.3 million cells completed within the hour (Figure 1).

**Figure 1.**
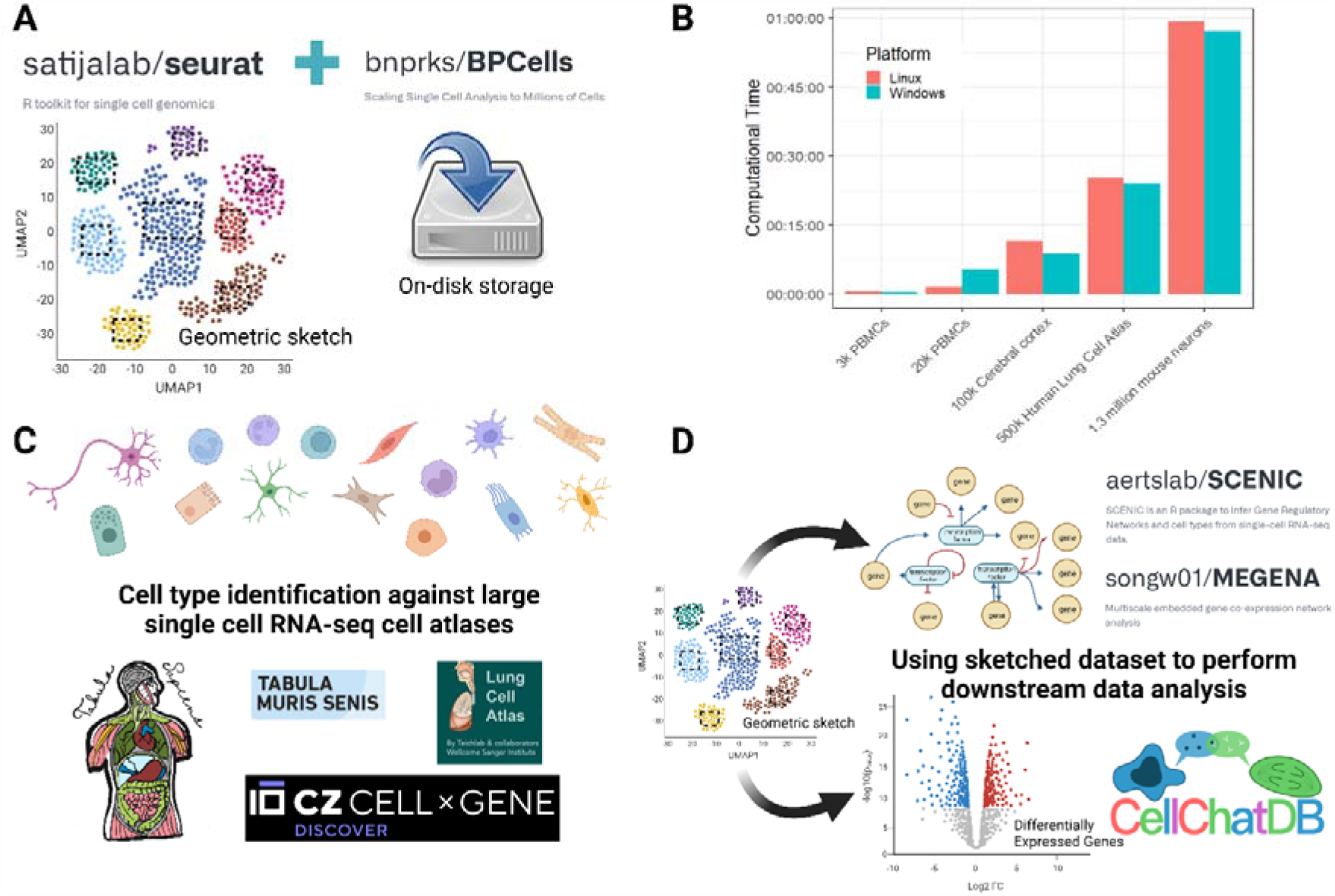
Efficient single cell RNA-seq analysis with ICARUS v3. (A) Efficient computational speed is achieved in ICARUS v3 through the use of a ‘geometric sketching’ method of sampling a subset of representative cells whilst the larger overall dataset is stored on-disk. Dimensionality reduction and clustering is performed on the sketched dataset and then projected back onto the overall dataset. (B) Benchmarking of ICARUS v3 dimensionality reduction and clustering for various datasets of increasing cell numbers (Dataset list available in Supplementary 1). A sketched dataset of 10,000 cells was taken. Each sketched dataset was scaled and log normalised and dimensionality reduction was performed using 2000 variable features (Seurat::FindVariableFeatures) and the first 50 PCA dimensions. Graph based clustering was performed using the Louvain algorithm with a k-nearest value of 20. Benchmarking was performed on a Linux Ubuntu 22.04.2 LTS with 64GB RAM and AMD EPYC-Milan Processor with 16 CPU cores. Benchmarking was also assessed on a Windows 11 machine with 16GB ram running a 16-core 11th Gen Intel(R) Core(TM) i7-11800H @2.30GHz. (C) ICARUS v3 introduces cell type annotation against large single cell atlases including Tabula sapiens, Tabula muris senis, human lung cell atlas and others publicly available in the Chan Zuckerberg CELLxGENE database. (D) The geometric sketched dataset is leveraged to perform common downstream data analysis including co-expression network analysis, gene regulatory network construction, trajectory analysis, cell-cell signalling and examination of cell cluster association with GWAS traits.

### Cell type annotation against single cell atlases

Another major update introduced in ICARUS v3 is the incorporation of large single cell RNA-seq atlases comprising of millions of cells for cell cluster labelling. ICARUS now supports cell label transfer utilising the SingleR method (10) for atlases including Tabula sapiens (2), Tabula muris sensis (1), Human Brain Cell Atlas v1.0 (11), Human Lung Cell Atlas (12), Asian Immune Diversity Atlas (AIDA) (https://chanzuckerberg.com/science/programs-resources/single-cell-biology/ancestry-networks/immune-cell-atlas-of-asian-populations/), developing human immune system (13), healthy human liver (14) and adult human retina (15). Furthermore, datasets from the Chan Zuckerberg CELLxGENE (CZ CELLxGENE) database may be directly loaded into ICARUS and cell type labels transferred using SingleR methodology. ICARUS additionally retains the functionality to perform cell cluster labelling through sctype, a R package that congregates cell type specific markers from CellMarker (http://biocc.hrbmu.edu.cn/CellMarker) and PanglaoDB (https://panglaodb.se) databases. To achieve efficient cell type annotation, the subset of sketched cells is first annotated against the reference datasets using SingleR or sctype and then projected back onto the larger overall dataset.

### Doublet detection at scale

We have previously introduced the DoubletFinder (16) methodology of identifying cell multiplets that may arise during single cell RNA-seq library generation. However, the computational speed of DoubletFinder during artificial k nearest neighbour (pANN) simulation is not efficient for large datasets (17). ICARUS v3 utilises the subset of sketched cells to perform pANN generation at a small scale which then are projected back to the overall dataset (ProjectData functionality of Seurat v5) to enable approximation of multiplets at a large scale.

### Streamlined incorporation of 10X Genomics and Anndata hdf5 files

We also introduce an easier method of data input with support for the 10X Genomics hdf5 and Anndata hdf5 file formats (18). Users may now upload multiple hdf5 files at once for streamlined integration. Integration of datasets may be performed using anchor-based CCA integration (19), anchor-based RPCA integration (19), harmony (20) or fastMNN (21).

## Summary

Our latest update to ICARUS provides users with the capability to process large datasets at speed that previously could not be effectively processed. To our knowledge, ICARUS is currently the only publicly accessible web server that supports in depth analysis of large-scale single cell RNA-seq data. Moreover, users can take advantage of ICARUS’s built-in save and load feature, which has also been updated to leverage on-disk storage to streamline analysis and minimize computational time spent for repeated analyses requiring resource-intensive steps. ICARUS will continue to receive ongoing updates as new methodologies are developed, ensuring that users have access to a cutting-edge resource for making novel discoveries.

## Data availability

The functionality of ICARUS v3 was demonstrated on 5 datasets of increasing cell numbers (Supplementary 1). Data is available from 10X Genomics and ChanZuckerberg CellxGene databases. Refer to Supplementary 1 for full details. The datasets are also fully accessible from ICARUS v3 application.

## Benchmarking

For benchmarking, a sketched dataset of 10,000 cells was generated using the SketchData function from Seurat whilst the large overall dataset was stored on disk using the BPCells write_matrix function. The sketched dataset was then scaled and log normalised and dimensionality reduction was performed using 2000 variable features (Seurat::FindVariableFeatures) and the first 50 PCA dimensions. Graph based clustering was performed using the Louvain algorithm with a k-nearest value of 20. Benchmarking was performed on a Linux Ubuntu 22.04.2 LTS with 64GB RAM and an AMD EPYC-Milan Processor with 16 CPU cores. Benchmarking was also assessed on a Windows 11 machine with 16GB ram running a 16-core 11th Gen Intel(R) Core(TM) i7-11800H @2.30GHz.

## Code availability

ICARUS is available at https://launch.icarus-scrnaseq.cloud.edu.au/. The application is free and open to all users with no login requirement. For data privacy reasons, the user data is not retained on the server after the user-session is terminated.

R source code of the ICARUS v3 shiny app is available at 10.5281/zenodo.10155798. Alternatively, a docker version is accessible through the Docker Hub under the name ‘icarusscrnaseq/icarus_v3’.

## Acknowledgements

We thank the New Zealand–China Non-Communicable Diseases Research Collaboration Centre (NCD CRCC). This web server is supported by the Australian National eResearch Collaboration Tools and Resources (Nectar Research Cloud) initiative.

## Funding

New Zealand Ministry of Business Innovation and Employment funding for New Zealand– China Non-Communicable Diseases Research [UOOX1601]. Funding for open access charge: New Zealand–China Non-Communicable Diseases Research [UOOX1601].

## Contributions

A.J., K.L., and R.S., considered the study. A.J., developed the method, implemented the webserver tool, and analysed the data. A.J., wrote the manuscript. A.J., K.L., and R.S., supervised the study, tested the webserver, and revised the manuscript. All the authors read and approved the final manuscript.

## Notes

### Competing Interest Statement

The authors have declared no competing interest.

### Summary of Updates

Link to web server in abstract was incorrect

