## Supplementary 1 for "ICARUS v3, a massively scalable web server for single cell RNA-seq analysis of millions of cells"

**Supplementary 1 Example single cell RNA-seq datasets used for benchmarking in ICARUS v3**

| **Dataset** | **Number of Cells** | **Available from…** |
| --- | --- | --- |
| 3k Human peripheral blood mononuclear cells (Seurat guided clustering tutorial dataset) | 3,156 | 10x Genomics  <https://cf.10xgenomics.com/samples/cell/pbmc3k/pbmc3k_filtered_gene_bc_matrices.tar.gz> |
| 20k Human peripheral blood mononuclear cells | 23,747 | 10x Genomics  <https://www.10xgenomics.com/resources/datasets/20-k-human-pbm-cs-3-ht-v-3-1-chromium-x-3-1-high-6-1-0> |
| 100k Cerebral cortex (Cx) - Precentral gyrus (PrCG) - Primary motor cortex - M1C from Human Brain Cell Atlas | 116,576 | CZ CellxGene  <https://cellxgene.cziscience.com/collections/283d65eb-dd53-496d-adb7-7570c7caa443> |
| 500k Human Lung Cell Atlas | 584,944 | CZ CellxGene  <https://cellxgene.cziscience.com/collections/6f6d381a-7701-4781-935c-db10d30de293> |
| 1.3 Million Brain Cells from E18 Mice | 1,306,127 | 10x Genomics  <https://www.10xgenomics.com/resources/datasets/1-3-million-brain-cells-from-e-18-mice-2-standard-1-3-0> |
